# Combating viral contaminants in CHO cells by engineering STAT1 mediated innate immunity

**DOI:** 10.1101/423590

**Authors:** Austin W.T. Chiang, Shangzhong Li, Benjamin P. Kellman, Gouri Chattopadhyay, Yaqin Zhang, Chih-Chung Kuo, Jahir M. Gutierrez, Faeazeh Ghazi, Hana Schmeisser, Patrice Ménard, Sara Petersen Bjørn, Bjørn G. Voldborg, Amy S. Rosenberg, Montserrat Puig, Nathan E. Lewis

## Abstract

Viral contamination in biopharmaceutical manufacturing can lead to shortages in the supply of critical therapeutics. To facilitate the protection of bioprocesses, we explored the basis for the susceptibility of CHO cells, the most commonly used cell line in biomanufacturing, to RNA virus infection. Upon infection with certain ssRNA and dsRNA viruses, CHO cells fail to generate a significant interferon (IFN) response. Nonetheless, the downstream machinery for generating IFN responses and its antiviral activity is intact in these cells: treatment of cells with exogenously-added type I IFN or poly I:C prior to infection limited the cytopathic effect from Vesicular stomatitis virus (VSV), Encephalomyocarditis virus (EMCV), and Reovirus-3 virus (Reo-3) in a STAT1-dependent manner. To harness the intrinsic antiviral mechanism, we used RNA-Seq to identify two upstream repressors of STAT1: Gfi1 and Trim24. By knocking out these genes, the engineered CHO cells exhibited increased resistance to the prototype RNA viruses tested. Thus, omics-guided engineering of mammalian cell culture can be deployed to increase safety in biotherapeutic protein production among many other biomedical applications.

## Introduction

Chinese hamster ovary (CHO) cells are extensively used to produce biopharmaceuticals^1^ for numerous reasons. Though one advantage is their reduced susceptibility to many human virus families^2-4^, there have been episodes of animal viral contamination of biopharmaceutical production runs, mostly from trace levels of viruses in raw materials. These infections have led to expensive decontamination efforts and threatened the supply of critical drugs^5-7^. Viruses that have halted production of valuable therapeutics include RNA viruses such as Cache Valley virus^6^, Epizootic hemorrhagic disease virus^8^, Reovirus^6^ and Vesivirus 2117^9^. Thus, there is a critical need to understand the mechanisms by which CHO cells are infected and how the cells can be universally engineered to enhance their viral resistance^10^. For example, a strategy was proposed to inhibit infection of CHO cells by minute virus of mice by engineering glycosylation^11^. We present an alternative strategy to prevent infections of a number of RNA viruses with different genomic structures and strategies to interfere with the host anti-viral defense.

Many studies have investigated the cellular response to diverse viruses in mammalian cells, and detailed the innate immune responses that are activated upon infection. For example, type I interferon (IFN) responses play an essential role in regulating the innate immune response and inhibiting viral infection^12-15^ and can be induced by treatment of cells with poly I:C^16-18^. However, the detailed mechanisms of virus infection and the antiviral response in CHO cells remain largely unknown. Understanding the role of type I IFN-mediated innate immune responses in CHO cells could be invaluable for developing effective virus-resistant CHO bioprocesses. Fortunately, the application of recent genome sequencing^19-23^ and RNA-Seq tools can now allow the analysis of complicated cellular processes in CHO cells^24-28^, such as virus infection.

To unravel the response of CHO cells to viral infection, we infected CHO-K1 cells with RNA viruses from diverse virus families. We have further assayed the ability of activators of type I IFN pathways to induce an antiviral response in the cells. Specifically, we asked the following questions: (1) Can CHO-K1 cells mount a robust type I IFN response when infected by RNA viruses? (2) Can innate immune modulators trigger a type I IFN response of CHO-K1 cells and, if so, are the type I IFN levels produced sufficient to protect CHO-K1 cells from RNA virus infections? (3) Which biological pathways and processes are activated during virus infection and/or treatment with innate immune modulators, and are there common upstream regulators that govern the antiviral response? (4) Upon the identification of common upstream regulators, how can we engineer virus resistance into CHO cells for mitigating risk in mammalian bioprocessing? Here we address these questions, illuminate antiviral mechanisms of CHO cells, and guide the development of bioprocess treatments and cell engineering efforts to make CHO cells more resistant to viral infection.

## Materials and Methods

### CHO-K1 cells and RNA virus infections

The susceptibility of CHO-K1 cells to viral infection has been previously reported^3^. Since infectivity was demonstrated for viruses of a variety of families (harboring distinct genomic structures), we selected the following RNA viruses from three different families to be used as prototypes: Vesicular stomatitis virus (VSV, ATCC^®^ VR-1238), Encephalomyocarditis virus (EMCV, ATCC^®^ VR-129B), and Reovirus-3 virus (Reo-3, ATCC^®^ VR-824). Viral stocks were generated in susceptible Vero cells as per standard practices using DMEM (Dulbecco’s Modified Eagle’s medium) supplemented with 10% FBS, 2mM L-glutamine, 100 U/ml penicillin and 100 μg/ml streptomycin (DMEM-10). Viral stocks were titered by tissue culture infectious dose 50 (TCID_50_) on CHO-K1 cells and used to calculate the multiplicity of infection in the experiments (Table 1).

**Table 1.** Study prototype viruses and multiplicity of infection (MOI) on CHO-K1 cells.

| Virus | Virus family | Genomic nucleic acid nature | Referenced CHO cell culture infection | MOI |
| --- | --- | --- | --- | --- |
| Vesicular stomatitis virus (VSV) | Rabdo­viridae | ss (-) RNA | Potts, 2008 | 0.003 |
| Encephalomyocarditis virus (EMCV) | Picornaviridae | ss (+) RNA | Potts, 2008 | 0.007 |
| Reovirus 3 (Reo-3) | Reoviridae | ds RNA | Wisher, 2005; Rabenau 1993 | 0.0013 |

#### Virus infection procedures

Cells were seeded in cell culture plates (3×10^5^ and 1.2×10^6^ cells/well in 96-well and 6-well plates, respectively) and grown overnight in RPMI-1040 supplemented with 10% FBS, 2mM L-glutamine, 100 U/ml penicillin and 100 μg/ml streptomycin, 10 mM Hepes, 1x non-essential amino acids and 1 mM sodium pyruvate (RPMI-10). IFNα/β (human IFNα (Roferon) and IFNβ (Avonex), mouse IFNα (Bei Resources, Manassas, VA)) as well as innate immune modulators (LPS (TLR4) (Calbiochem), CpG-oligodeoxynucleotide (ODN) D-ODN, 5’-GGTGCATCGATGCAGGGGG-3’^29^ and ODN-1555, 5’-GCTAGACGTTAGCGT-3’ (TLR9) (custom-synthesized at the Center for Biologics Evaluation and Research facility, FDA), imidazoquinoline R837 (TLR7/8) (Sigma) and poly I:C-Low molecular weight/LyoVec (poly I:C) (Invivogen) were added to the cultures 24 h prior to testing or virus infection, at the concentrations indicated in the figures. Note that, by monitoring changes in the gene expression levels of IFNβ and Mx1 in the cells, we established that 16-20 h would be an adequate time interval for treating cells with poly I:C prior to infection (Figure S1). Anti-IFNβ neutralizing antibody (2.5 μg/ml; Abcam, Cambridge, MA cat# 186669) was also used in certain experiments, 24 h prior to infection. Viral infection was performed by adding virus suspensions to the cell monolayers at the indicated MOI in serum-free media and incubated at 37 °C, 5% CO_2_ for 2h. Cell cultures were washed twice to discard unbound virus and further incubated at 37 °C for 30 h (VSV), 54 h (EMCV) or 78 h (Reo-3) (unless otherwise indicated in the figures). The cell harvesting time was established based on appearance of cytopathic effect in approximately 50% of the cell monolayer. Cytopathic effect was visualized by crystal violet staining as per standard practices. Infection/poly I:C experiments were repeated twice, independently. In each experiment, CHO cells were cultured as poly I:C untreated – uninfected (media control, m), poly I:C treated – uninfected (p), poly I:C untreated – virus infected (Vm) and poly I:C treated – virus infected (Vp).

#### Western blot procedures

Cell lysates were prepared using mammalian protein extraction reagent M-PER (Thermo Fisher Scientific, Waltham, MA) with Protease and Halt^™^ phosphatase inhibitor cocktails (Thermo Fisher Scientific) using an equal number of cells per sample. Samples were analyzed by SDS-PAGE using 10-20% Tris-Glycine gels (Thermo Fisher Scientific) under reducing conditions.

As a molecular weight marker, protein ladder (cat# 7727S) from Cell Signaling Technology (Danvers, MA) was used. Nitrocellulose membranes and iBlot^™^ transfer system (Thermo Fisher Scientific) were used for Western Blot analysis. All other reagents for Western Blot analyses were purchased from Thermo Fisher Scientific. Membranes were blocked with nonfat dry milk (BIO-RAD, Hercules, CA) for 1h followed by incubation with primary antibodies against STAT1, pSTAT1 (pY701, BD Transduction Lab, San Jose, CA), pSTAT2 (pY689, Millipore Sigma, Burlington, MA), actin (Santa Cruz Biotechnology, Santa Cruz, CA) or Mx1 (gift from O. Haller, University of Freiburg, Freiburg, Germany) O/N at 4°C. Secondary goat anti-mouse and anti-rabbit antibodies were purchased from Santa Cruz Biotechnology. SuperSignal West Femto Maximum Sensitivity Kit (Thermo Fisher Scientific) was used to develop membranes, and images were taken using LAS-3000 Imaging system (GE Healthcare Bio-Sciences, Pittsburgh, PA).

### RNA extraction, purification, and real-time PCR

Cell cultures were re-suspended in RLT buffer (Qiagen) and kept at -80°C until RNA was extracted using the RNeasy kit (Qiagen) and on-column DNAse digestion. RNA was eluted in 25 μl of DEPC water (RNAse/DNAse free); concentration and purity were tested by bioanalyzer. Total RNA levels for type I IFN related genes and viral genome were also assessed by RT-PCR. Complementary DNA synthesis was obtained from 1 μg of RNA using the High capacity cDNA RT kit (Thermo Fisher scientific) as per manufacturer’s instructions. Semi-quantitative PCR reactions (25 μl) consisted in 1/20 cDNA reaction volume, 1x Power Sybr master mix (Thermo Fisher Scientific), 0.5 μM Chinese hamster-specific primers for IFNβ, Mx1, IRF7 and IITMP3 sequences (SAbiosciences). Eukaryotic 18S was used as a housekeeping gene and assessed in 1X Universal master mix, 18S expression assay (1:20) (Applied Biosystems) using a 1/50 cDNA reaction volume. Fold changes were calculated by the 2-ΔΔCt method.

### cDNA library construction and Next-generation sequencing (RNA-Seq)

Library preparation was performed with Illumina’s TruSeq Stranded mRNA Library Prep Kit High Throughput (Catalog ID: RS-122-2103), according to manufacturer’s protocol. Final RNA libraries were first quantified by Qubit HS and then QC on Fragment Analyzer (from Advanced Analytical). Final pool of libraries was run on the NextSeq platform with high output flow cell configuration (NextSeq^®^ 500/550 High Output Kit v2 (300 cycles) FC-404-2004). Raw data are deposited at the Gene Expression Omnibus and Short Read Archive (accession numbers: GSE119379)

### RNA-Seq quantification and differential gene expression analysis

RNA-Seq quality was assessed using FastQC. Adapter sequences and low-quality bases were trimmed using Trimmomatic^30^. Sequence alignment was accomplished using STAR^31^ against the CHO genome (GCF_000419365.1_C_griseus_v1.0) with default parameters. HTSeq^32^ was used to quantify the expression of each gene. We performed differential gene expression analysis using DESeq2^33^. After Benjamini-Hochberg FDR correction, genes with adjusted p-values less than 0.05 and fold change greater than 1.5 were considered as differentially expressed genes (DEGs). Table S1 shows the number of identified DEGs in the three different comparisons: 1) untreated – uninfected vs. untreated – virus infected (m vs. Vm); 2) untreated – uninfected vs. poly I:C treated – uninfected (m vs. p); and 3) untreated – virus infected vs. poly I:C treated – virus infected (Vm vs. Vp).

### Genetic engineering (Gfi1, Trim24, Gfi1/Trim24) of CHO-S cell lines

CHO-S cells (Thermo Fisher Scientific Cat. # A1155701) and KO clones were cultured in CD CHO medium supplemented with 8 mM L-glutamine and 2 mL/L of anti-clumping agent (CHO medium) in an incubator at 37°C, 5% CO_2_, 95% humidity. Cells were transfected using FuGENE HD reagent (Promega Cat. # E2311). The day prior to transfection, viable cell density was adjusted to 8×10^5^ cells/mL in an MD6 plate well containing 3 mL CD CHO medium supplemented with 8 mM L-glutamine. For each transfection, 1500 ng Cas9-2A-GFP plasmid and 1500 ng gRNA plasmid (see Text S1 for details about the construction of plasmids) were diluted in 75 uL OptiPro SFM. Separately, 9 uL FuGene HD reagent was diluted in 66 uL OptiPro SFM. The diluted plasmid was added to the diluted FuGENE HD and incubated at room temperature for 5 minutes and the resultant 150 μL DNA/lipid mixture was added dropwise to the cells. For viability experiments, CHO-S KO cell lines were seeded at 3×10^6^ cells in 30 ml in CHO medium and incubated at 37 °C, 5% CO_2_, 125 rpm for up to 7 days. Infections were conducted with EMCV and Reo-3 at the same MOI calculated in CHO-K1 cells for 2h prior to wash cells twice to discard unbound particles. Control cell lines showing susceptibility to either virus were infected in parallel to those with Gfi1 and Trim24 gene KO.

## Results and Discussion

### CHO-K1 cells fail to resolve infection by RNA viruses despite possessing functional type I IFN-inducible anti-viral mechanisms

To evaluate the response of CHO cells to three different RNA viruses (VSV, EMCV and Reo-3; see Table 1), cells were infected and monitored for cytopathic effects and gene expression changes related to the type I IFN response. All three viruses induced a cytopathic effect (Figure 1A, right panels) and a modest increase in IFNβ transcript levels in infected CHO cell cultures was measured (Figure 1B), suggesting limited production of IFN. Through its cellular receptor, IFNα/β can further activate downstream interferon-stimulated genes known to limit viral infection both in cell culture and *in vivo*^34-37^. We noted that CHO cells seem to have a functional IFNα/β receptor and its activation with exogenous IFN confers resistance of CHO cells to VSV infection (see Supplementary Information Text S2 and Figure S2). Interestingly, CHO cells expressed high levels of the antiviral gene Mx1 when infected with Reo-3, but not VSV and EMCV (Figure 1C). Nevertheless, the virus-induced IFN mRNA response in the host cell was insufficient to prevent cell culture destruction. These data suggest a possible inhibition of the antiviral type I IFN response that varies across viruses, as previously reported^38-41^.

**Figure 1.**
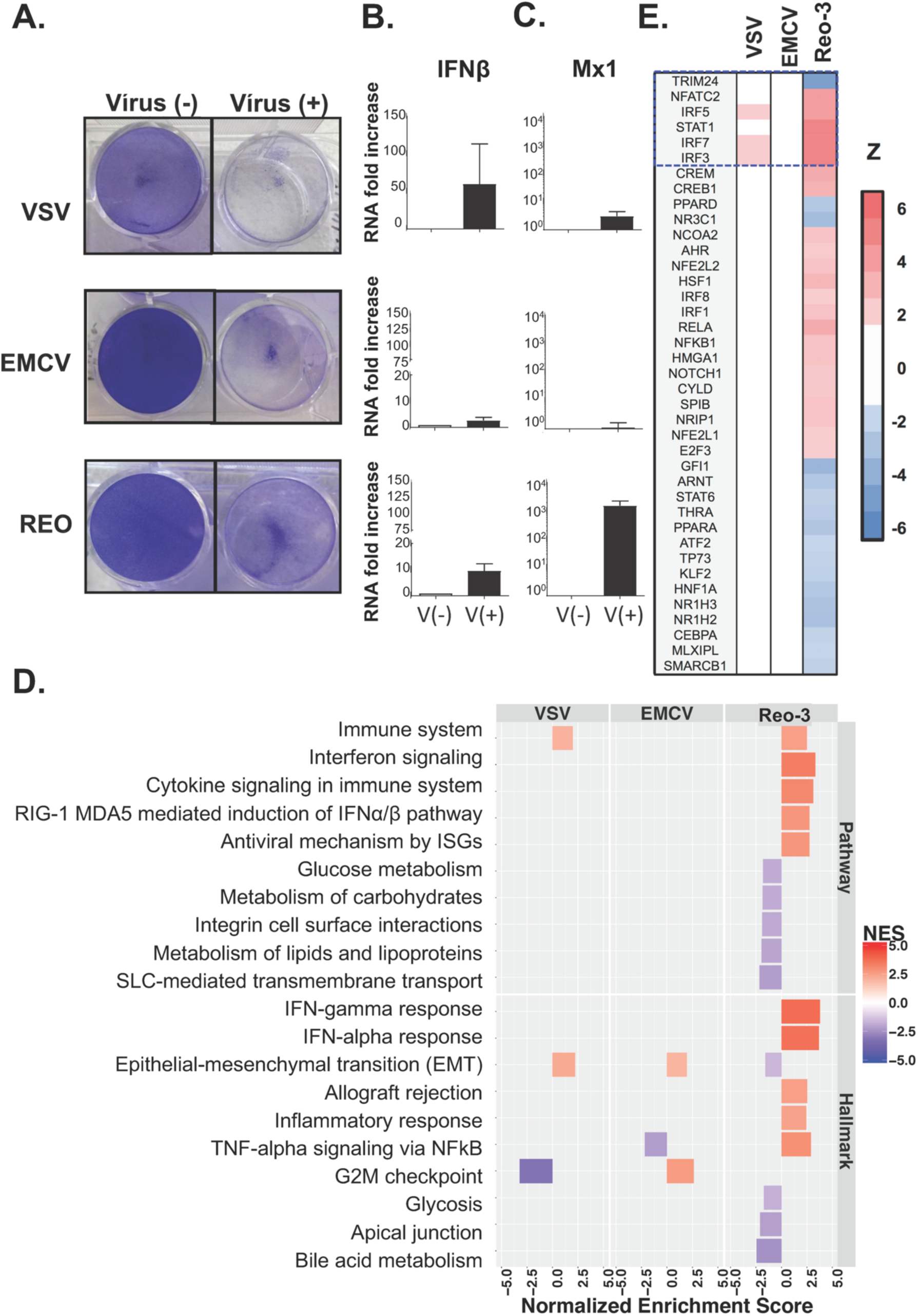
RNA viruses induce cytopathic effects on CHO-K1 cells. (A) Cytopathic effect of the three RNA viruses on CHO cells upon 30h (VSV), 54h (EMCV) or 78h (Reo-3) of infection. Fold change in IFNβ (B) and Mx1 (C) gene expressions in CHO cells infected with the three RNA viruses compared to uninfected cells at the same time points. (D) Several pathways and processes were enriched for differentially expressed genes following viral infection (m vs. Vm). (E) Top activated (red) or repressed (blue) upstream regulators following virus infection.

To explore why the induced type I IFN failed to mount a productive antiviral response in CHO cells, we conducted RNA-Seq and pathway analysis using GSEA (see details in Text S3 and Table S1). GSEA analysis that compared control vs. infected CHO cells (m vs. Vm) revealed the modulation of several immune-related gene sets and pathways activated by the virus (Figures 1D and S3, Table S2, and Text S4). Unlike VSV and EMCV, Reo-3 induced the ‘interferon alpha response’ and ‘RIG-I and MDA5-mediated induction of IFNα’ pathways ((p-value, NES) = (9.05×10^-3^, 3.68) and (1.12×10^-2^, 2.74), respectively). These findings were consistent with observations that the reovirus genome (dsRNA) can stimulate TLR3 and RIG-I to induce innate immune responses in other cell types^42-44^, in which the observed responses diverged markedly from the VSV and EMCV infections.

As we observed for Mx1, only Reo-3-infected cells showed a significant enrichment of differentially expressed genes involved in the type I IFN response (FDR-adjusted p-value = 9.05×10^-3^; normalized enrichment score, NES = 3.68). These genes contain the consensus transcription factor binding sites in the promoters that are mainly regulated by the transcription factor STAT1 and the interferon regulatory factors (IRF) family, such as IRF1, IRF3, IRF7 and IRF8 (Figure 1E). These results are consistent with observations that the IRF family transcription factors activate downstream immune responses in virus-infected mammalian cells^45, 46^. In contrast, VSV and EMCV failed to trigger anti-viral related mechanisms (e.g., type I IFN responses) downstream of IFNβ (Figures 1D and S3A). Examples of a few pathways that were stimulated included ‘immune system’ (including adaptive/innate immune system and cytokine signaling in immune system) in VSV (FDR-adjusted p-value = 1.49×10^-2^; normalized enrichment score, NES= 1.99) and the ‘G2M checkpoint’ in EMCV (p-value = 8.95×10^-3^; NES = 2.64). Disruption of the cell cycle affecting the G2M DNA checkpoint networkhas been reported for the survival of several viruses, including HIV (ssRNA)^47^, EBV (dsDNA)^48^, JCV (DNA)^49^, HSV (DNA)^50^. However, further studies will need to confirm whether VSV or EMCV use a similar strategy to escape the cell defense. Nevertheless, neither VSV nor EMCV infection activated known upstream activators of type I IFN pathways (Figure 1E) when analyzed with Ingenuity Pathway Analysis (IPA)^51^.

### Poly I:C induces a robust type I interferon response in CHO cells

Type I IFN responses limit viral infection^12-15^, and innate immune modulators^52-54^ mimic pathogenic signals and stimulate pattern recognition receptors (PRRs), leading to the activation of downstream immune-related pathways. Intracellular PRRs, including toll-like receptors (TLR) 7, 8 and 9, and cytosolic receptors RIG-I or MDA5, can sense viral nucleic acids and trigger the production of type I IFN. Thus, we asked whether CHO cell viral resistance could be improved by innate immune modulators.

CHO PRRs have not been studied extensively, so we first assessed the ability of synthetic ligands to stimulate their cognate receptors to induce a type I IFN response. CHO cells were incubated with LPS (TLR4 ligand), CpG-oligodeoxynucleotide (ODN) type D (activates TLR9 on human cells), ODN-1555 (activates TLR9 on murine cells), imidazoquinoline R837 (TLR7/8 ligand) and poly I:C-Low molecular weight/LyoVec (poly I:C) (activates the RIG-I/MDA-5 pathway), and subsequently tested for changes in expression of IFN stimulated genes with anti-viral properties. After 24 h of culture, gene expression levels of IRF7 and Mx1 increased significantly in cells treated with poly I:C but not in those treated with any of the other innate immune modulators (Figure 2A). Furthermore, STAT1 and STAT2 phosphorylation and Mx1 protein levels were elevated following treatment with poly I:C or exogenous interferon-alpha (IFNα), which was used as a control (Figure 2B and 2C).

**Figure 2.**
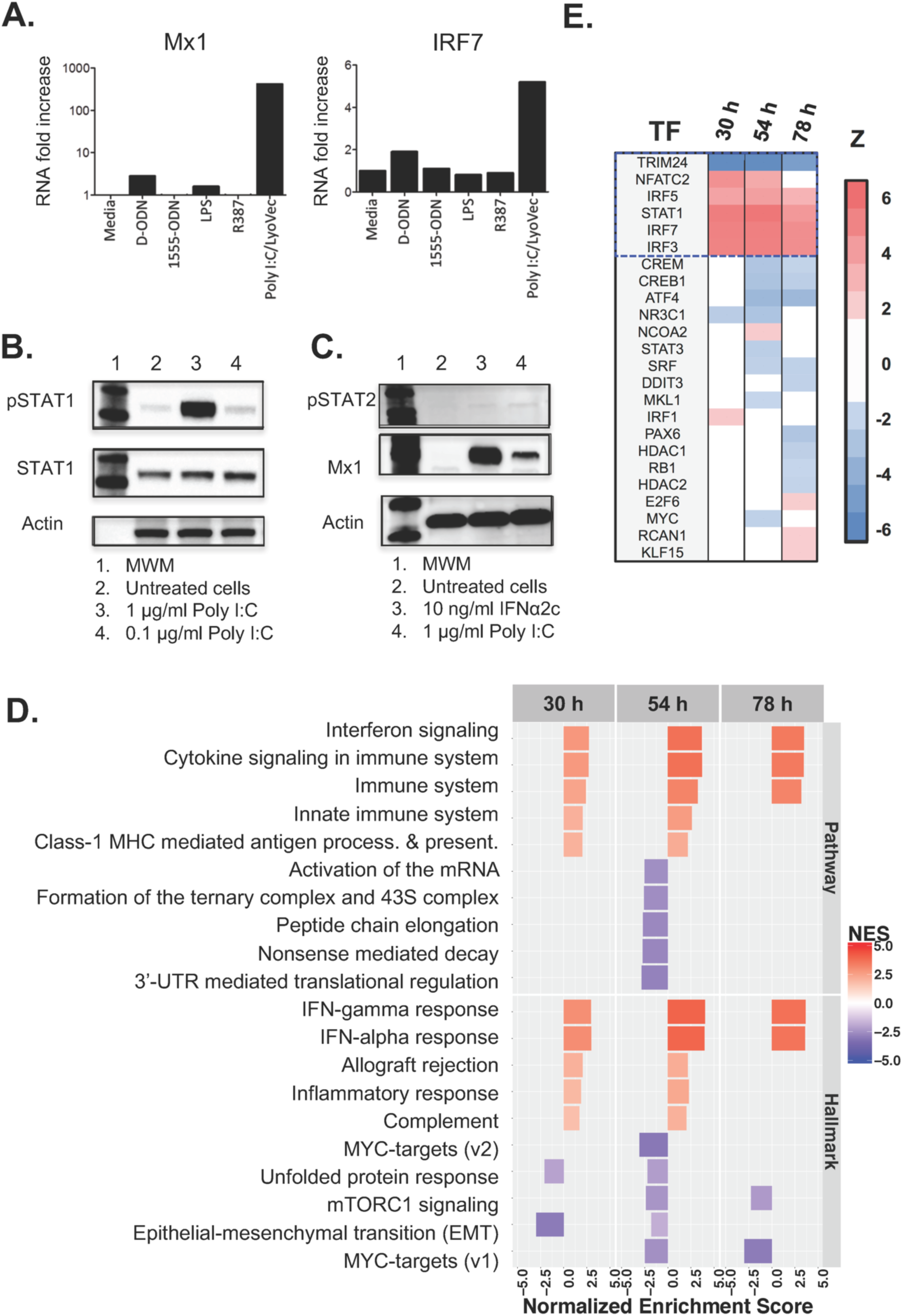
Innate immunity genes in CHO cells are activated by poly I:C. (A) IFN-stimulated transcription was increased in cells treated with poly I:C /LyoVec for 24h, but not with other TLR ligands engaging TLR9, TLR4 or TLR7/8. (B) Poly I:C triggered STAT1 phosphorylation in a dose dependent manner, and (C) the levels of STAT2 phosphorylation and Mx1 protein expression were comparable to those triggered by IFNα2c. (D) Several pathways and processes were enriched for differentially expressed genes following poly I:C treatment (m vs. p). (E) Top upstream regulators that are activated (red) or repressed (blue) following poly I:C treatment.

Next, we characterized the type I IFN response induced by poly I:C by analyzing the transcriptome of untreated vs. treated CHO cells. Cells were cultured with poly I:C in the media for 30, 54 and 78 h after an initial 16 h pre-incubation period (see Methods for details). GSEA of the RNA-Seq data demonstrated that poly I:C induced a strong ‘innate immune response’ in comparison to untreated cultures (media) (m vs. p; (p-value, NES, Enrichment strength) = (8.08×10^-3^, 2.98, 73%), (1.57×10^-2^, 3.95, 70%) and (3.91×10^-3^, 3.58, 78%)) evident in the three independently tested time points (Figures 2D and S3B, Text S4 and Table S3). In addition, poly I:C activated several upstream regulators of the type I IFN pathways (Figure 2E). We note that the GSEA strength (see Text S3) of the innate immune response induced by poly I:C (m vs. p) was stronger than the innate immune response seen for Reo-3 infection alone (m vs. Vm in Figure S3). Thus, CHO cells can activate the type I IFN signaling (JAK-STAT) pathway in response to poly I:C and display an anti-viral gene signature, which was sustained for at least 4 days.

### Poly I:C-induced type I interferon response protects CHO cells from RNA virus infections

We next examined if the type I IFN response, induced by poly I:C, could protect CHO cells from RNA virus infections. We found that poly I:C pre-treatment protected CHO cells against VSV infection through the IFNβ-mediated pathway (Figure S4 and Text S5), and that poly I:C protected against all three viruses tested (Figures 3A-C). Cell morphology differed notably between cultures infected with virus (Vm), control uninfected cells (m), and poly I:C pre-treated cultures (p and Vp) (Figures 3A-C, left panels). These morphological changes correlated with the cytopathic effect observed in the cell monolayers (Figures 3A-C, right panels). At 78h, the extent of cell culture damage by Reo-3, however, was milder than by VSV and EMCV at a shorter incubation times (30h and 54h, respectively) (Panels Vm in Figures 3A-C), possibly since Reo-3 induced higher levels of anti-viral related genes in the CHO cells but VSV and EMCV did not (Figures 1C, 1D and 1E). Notably, although poly I:C pre-treatment conferred protection of CHO cells to all three viral infections (Panels Vp in the Figure 3A-C), striking transcriptomic differences were observed (Table S4). Poly I:C pre-treatment significantly activated immune-related pathways and up-regulated type I IFN-related gene expression in CHO cells infected with VSV and EMCV when compared to non-poly I:C pre-treated cells that were infected (Vm vs. Vp) (Figures 3D-E, S5A-B and Table S5). Poly I:C pre-treatment was sufficient to induce a protective type I IFN response to VSV and EMCV. In contrast, for Reo-3 infection, pre-treatment with poly I:C did not further increase the levels of expression of IFN associated genes already observed in no pre-treated cells. The lack of enhanced expression of antiviral genes in Reo-3 Vm vs. Vp observed in the GSEA was further confirmed by Taqman analysis. A similar level of expression of anti-viral Mx1 and IITMP3 genes^55-58^ was obtained for CHO cells independently infected with Reo-3 (Vm), treated with poly I:C (p), or pre-treated with poly I:C and infected (Vp), which resulted in no differences in transcript levels when we compared Vm vs. Vp (Figure S5C). Nevertheless, the outcome of infection was surprisingly different in Vm or Vp samples. To understand these differences, we searched for genes that were differently modulated by poly I:C treatment in the context of Reo-3 infection. Indeed, we identified 30 genes (Figure S6 and Table S6) that were significantly up regulated (adjusted p-value <0.05, fold change >1.5) in the comparisons of m vs. Vp and m vs. p but not in the comparison of m vs. Vm. These genes are significantly enriched in 11 KEGG pathways related to host-immune response (e.g., antigen processing and presentation, p-value=3.4×10^-3^) and processes important to virus infection (e.g., endocytosis, p-value=2.5×10^-2^). We also observed many of these genes significantly enriched molecular functions: 1) RNA polymerase II transcription factor activity (11 genes; GO:0000981 FDR-adjusted p-value < 1.30×10^-15^) and 2) nucleic acid binding transcription factor activity (12 genes GO:0001071 FDR-adjusted p-value < 3.54×10^-15^) by gene set enrichment analysis (see Text S3 and Table S7). This suggests that poly I:C treatment, 16 hours prior to virus infection, pre-disposes the cell to adopt an antiviral state and might restore the host transcription machinery subverted by Reo-3 virus resulting in the protection of the CHO cells.

**Figure 3.**
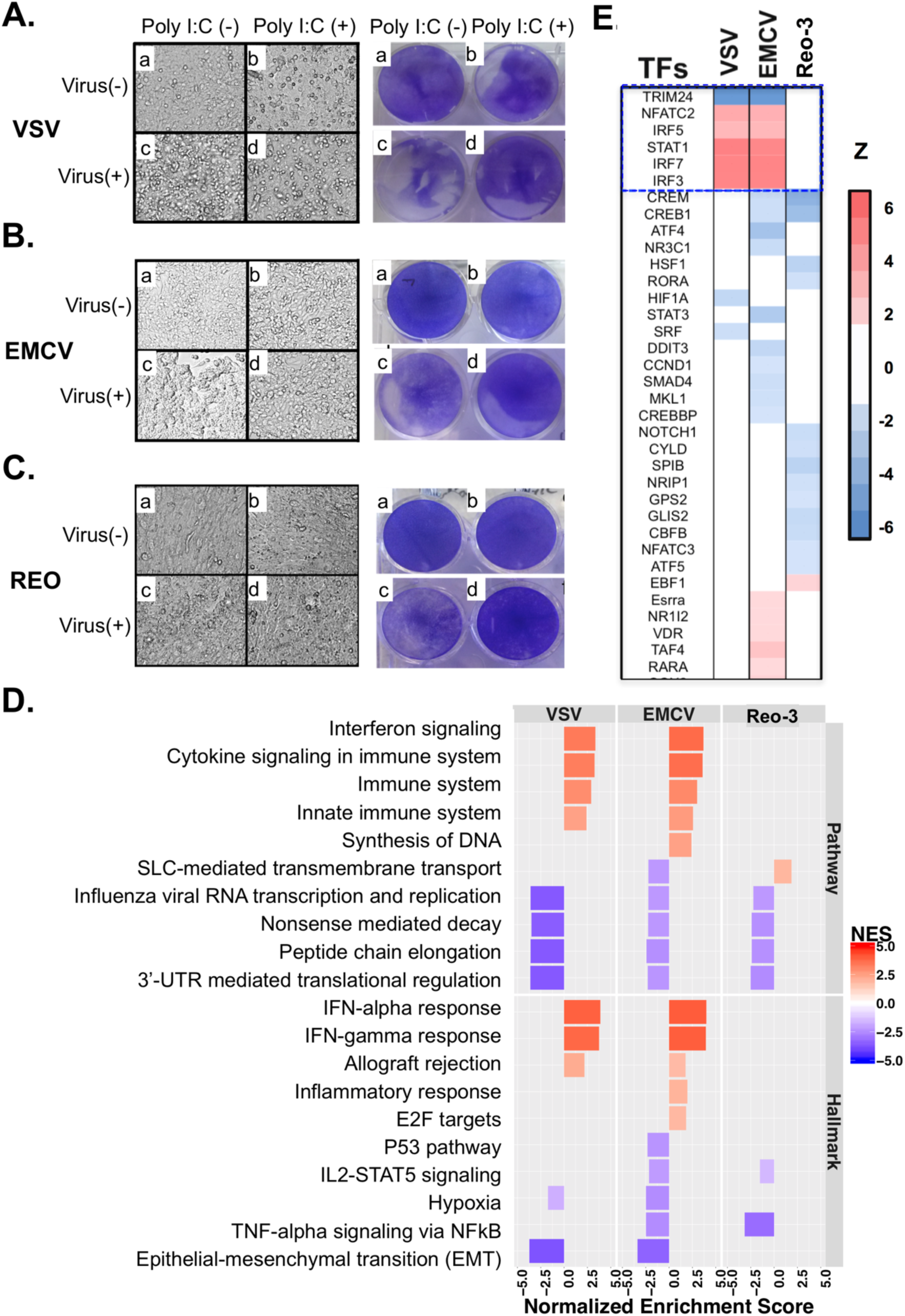
Poly I:C pre-treatment prevents virus infection of VCV, EMCV, and Reo-3. (A-C) Cell morphology (left panels) and cytopathic effect measured by crystal violet staining (right panels) of virus-infected CHO cells; (D) The enriched down-stream pathways under condition of Vm vs. Vp using RNA-Seq data. (E) The top 35 upstream regulators that are activated or repressed by poly I:C pre-treatment. A full list of the activated or repressed upstream regulators is shown in the Table S5.

Our results revealed other processes that are differentially activated or repressed between Vm and Vp (Figure 3D and Table S4). For example, the top down-regulated Reactome pathways in the virus-infected cells (Vm vs. Vp) are protein translational related processes: ‘nonsense mediated decay enhanced by the exon junction complex’ (p-value = 3.32×10^-2^, NES = -3.50), ‘peptide chain elongation’ (p-value = 3.32×10^-2^, NES = -3.59), and ‘3’-UTR mediated translational regulation’ (p-value = 3.38×10^-2^, NES = -3.61). These results agree with studies showing viral hijacking of the host protein translation machinery during infection^59^, and that the activation of interferon-stimulated genes restrain virus infections by inhibiting viral transcription and/or translation^12^. All these results suggest that poly I:C treatment provides the cell with an advantageous immune state that counteracts viral escape mechanisms and results in cell survival.

### A STAT1-dependent regulatory network governs viral resistance in CHO cells

GSEA revealed that several transcriptional regulators were activated or repressed during different viral infections and poly I:C-treated cells (Figures 1E, 2E, and 3E). Among these, NFATC2, STAT1, IRF3, IRF5, and IRF7 were consistently activated by poly I:C pre-treatment of CHO cells (m vs. p and Vm vs. Vp), and TRIM24 was suppressed. These transcription factors are involved in TLR-signaling (IRF3, IRF5, and IRF7)^45^ and JAK/STAT signaling (NFATC2, STAT1, and TRIM24). The TLR signaling pathway is a downstream mediator in virus recognition/response and in activating downstream type-I interferon immune responses^60-62^. Meanwhile, the JAK/STAT pathway contributes to the antiviral responses by up-regulating interferon simulated genes to rapidly eliminate virus within infected cells^63-65^. Importantly, one mechanism by which STAT1 expression and activity may be enhanced is via the poly I:C-induced repression of TRIM24 (an inhibitor of STAT1). The crosstalk between TLR- and JAK/STAT-signaling pathways is therefore important in virus clearance of infected host cells^66^.

In order to better understand the role of upstream regulators in the CHO cell viral protection, we examined the expression of the affected downstream target genes. Table 2 shows the regulatory pathways modulated by poly I:C treatment in uninfected (m vs. p; Table 2A) or infected (Vm vs. Vp; Table 2B) cells, and the described downstream effect. In cells surviving VSV and EMCV infection (Vp), we identified regulatory networks involved in restricting viral replication (Table 2B and Figures 4A and 4B). These networks are predominantly regulated by the 6 transcription factors (NFATC2, STAT1, IRF3, IRF5, IRF7, and TRIM24) that were also identified as transcription factors induced in poly I:C treated uninfected cells (p) (Table 2A). These findings suggest that the induction of the STAT1-dependent regulatory network by poly I:C treatment allows the cell to adopt an activated state that makes it refractory to virus infection. In contrast, the STAT1-dependent regulatory network was not apparent when comparing Reo-3 infected cells untreated and treated with poly I:C (Vm vs. Vp), because both Reo-3 and poly I:C induce STAT1 in CHO cells (Figure 1E and 2E). Poly I:C is a structural analog of double-stranded RNA and activates similar pathways as Reo-3^67^, such as, the NFATC2-dependent (Figures S7) and IRF3-dependent networks (Figures S8).

**Table 2A.** The downstream effects of the upstream regulators from the comparison of m vs. p.

| Virus | Consistency score<sup>*a</sup> | Total nodes (TF, TG, BP) | Transcription factors (TF)<sup>*b</sup> | Target gene (TG)<sup>*c</sup> | Biological Process (BP)<sup>*d</sup> | Relations<sup>*e</sup> |
| --- | --- | --- | --- | --- | --- | --- |
| 30h | 5.82 | 21<br>(5, 13, 3) | STAT1, IRF3, IRF5, IRF7, NFATC2 | CASP1, CXCL10, DDX58, EIF2AK2, IFIH1, IL15, ISG15, Mx1/Mx2, OASL2, PELI1, PML, SOCS1, TNFSF10 | **<u>Inhibit</u>**<br>Replication of virus.<br>**<u>Activate</u>**<br>Activation of phagocytes; Apoptosis of antigen presenting cells. | 6/15<br>(40%) |
| 54 h | 22.47 | 48<br>(7.29.12) | STAT1, IRF3, IRF5, IRF7, NFATC2, TRIM24, NCOA2 | BST2, C3, CASP1, CXCL10, DDX58, EGR2, EIF2AK2, GBP2, IFIH1, IFIT1B, IFIT2, IFITM3 (IITMP3), Igtp, IL15, ISG15, Mx1/Mx2, MYC, OASL2, PML, PSMB10, PSMB8, PSME2, PTGS2, SPP1, STAT2, TAP1, TLR3, TNFSF10, TRAFD1 | **<u>Inhibit</u>**<br>Replication of virus; Infection by RNA virus; Infection of central nervous system.<br>**<u>Activate</u>**<br>Antiviral response; Clearance of virus; Immune response of antigen presenting cells; Immune response of phagocytes; Cytotoxicity of leukocytes; Function of leukocytes; Infiltration by T lymphocytes; Quantity of MHC Class I of cell surface; Cell death of myeloid cells. | 21/84<br>(25%) |
| 78 h | 27.80 | 30<br>(8, 14, 8) | STAT1, IRF5, NFATC2, NR3C1, PPARD, ZBTB16, CDKN2A, EBF1 | C3, CCL2, CCL7, CD36, CXCL10, CXCL9, DDX58, EIF2AK2, ISG15, MYC, THBS1, TLR3, TNFSF10, VEGFA | **<u>Activate</u>**<br>Activation of macrophages; Apoptosis of myeloid cells; Cell movement of T lymphocytes; Cellular infiltration by leukocytes; Damage of lung; Recruitment of leukocytes; Response of myeloid cells; Response of phagocytes. | 11/64<br>(17%) |
| 78 h | 7.56 | 12<br>(2, 7, 3) | CDKN2A, ZBTB16 | C3, CCL2, CCL7, CXCL10, CXCL9, MYC, VEGFA | **<u>Activate</u>**<br>Cell movement of T lymphocytes; Recruitment of leukocytes; Survival of organism. | 1/6<br>(17%) |

**Table 2B.**
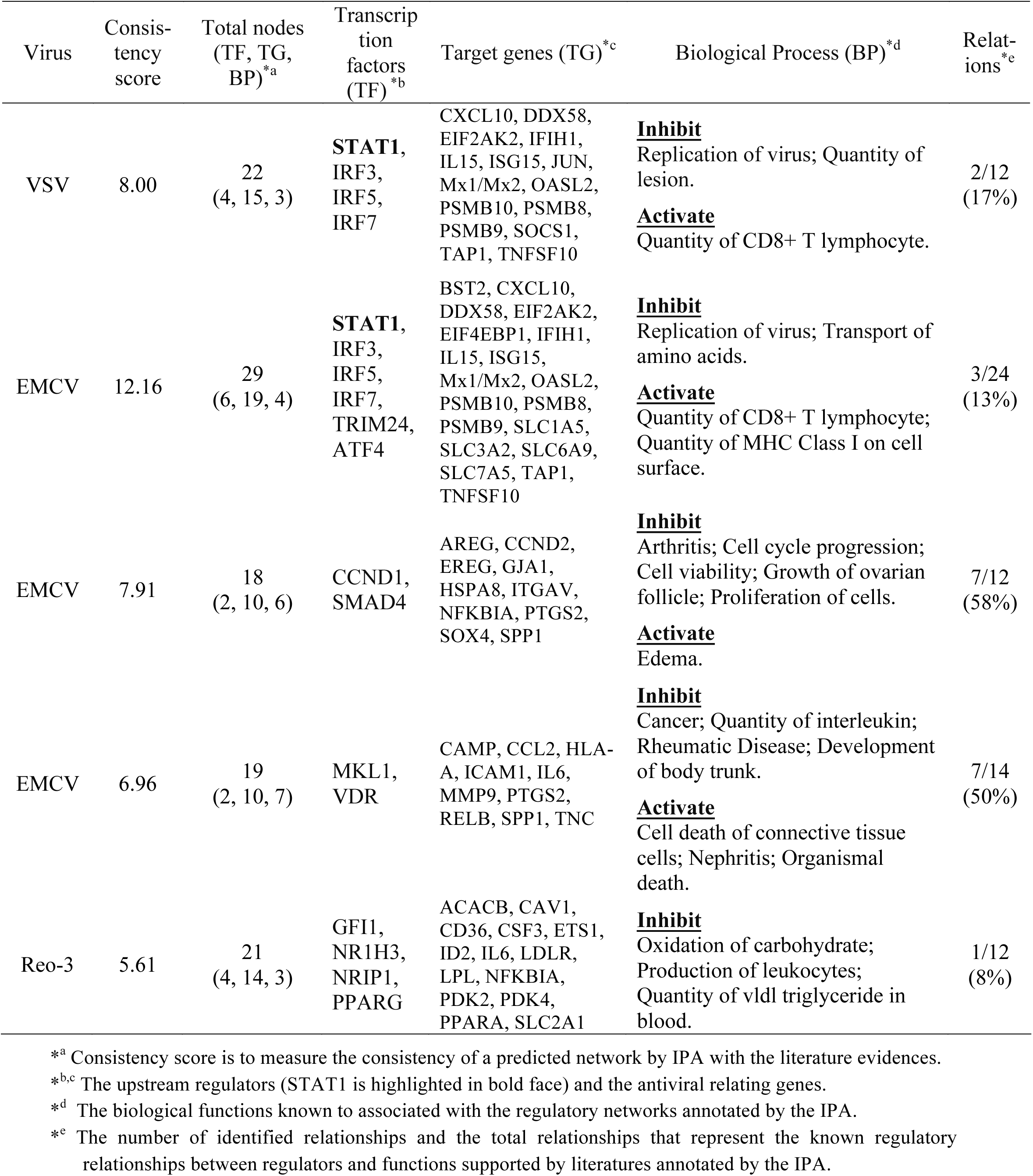
The downstream effects of the upstream regulators from the comparison of Vm vs. Vp.

**Figure 4.**
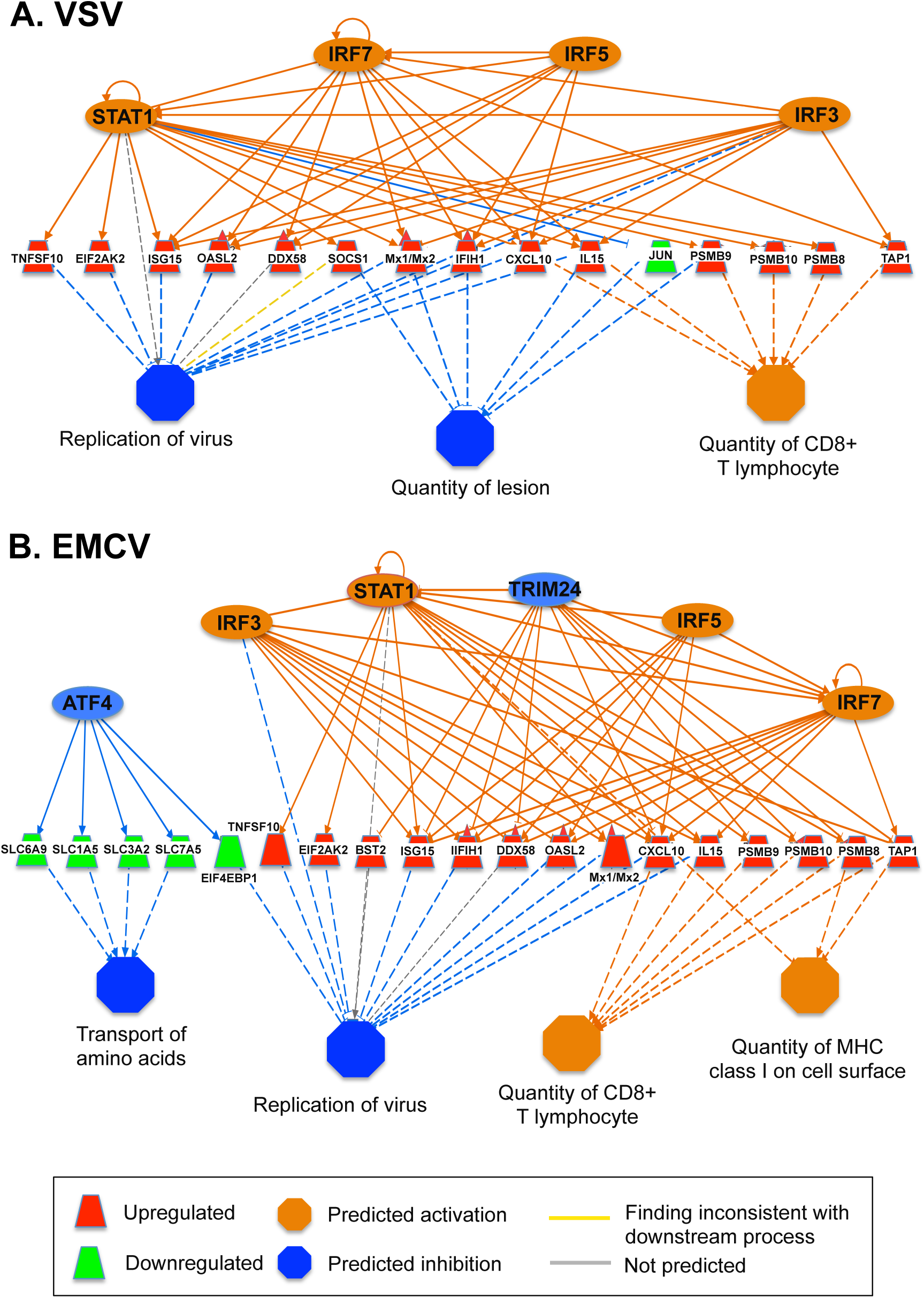
A STAT1-dependent regulatory network controls viral resistance (VSV and EMCV) in CHO cells. A STATl-dependent regulatory network induced by the pre-treatment of poly I:C leads to the inhibition of VSV (A) and EMCV (B) replication in CHO cells, based on the comparison of Vm and Vp RNA-Seq. The colors denote the states inferred from the RNA-Seq data. For example, the blue color of TRIM24 means that TRIM24 activity is suppressed, based on the differential expression of genes that are regulated by TRIM24.

With the STAT1 network potentially contributing to viral resistance, we searched for upstream regulators that could be modulated to maximally induce STAT1. We identified sixteen statistically significant (p < 0.05) upstream regulators, including 13 positive and 3 negative regulators of STAT1 using IPA (Figure 5; see details in Text S6). We hypothesized that the deletion of the most active repressors of STAT1 could improve virus resistance by inducing STAT1 gene expression and the downstream type I IFN antiviral response in the cell (Figure 5). We identified three STAT1 repressors (Trim24, Gfi1 and Cbl) with a negative regulatory score and therefore potential for inhibiting STAT1 based on the RNA-Seq differential expression data (see details in Text S6 and Figure S9). However, Cbl was not present in cells infected with Reo-3 (Table S8). Therefore, we selected the two negative regulators, Gfi1^68^ and Trim24^69^ of STAT1 as knockout targets for genetic engineering in CHO-S cells and subsequently tested the virus susceptibility of such KO cells, using Reo-3 and EMCV. We found that the Trim24 and Gfi1 single knockout clones showed resistance to Reo-3 but moderate or no resistance against EMCV (Figure 6A-B), compared to virus susceptible positive control cell lines (Figure S10). However, the Gfi1 + Trim24 double knockout (Figure 6C) showed resistance to both viruses tested, even when cells were passaged and cultured for an additional week (Figure S11). Together these results show that eliminating repressors of the STAT1 regulatory network contributes to antiviral mechanisms of CHO cells, which could possibly be harnessed to obtain virus-resistant CHO bioprocesses.

**Figure 5.**
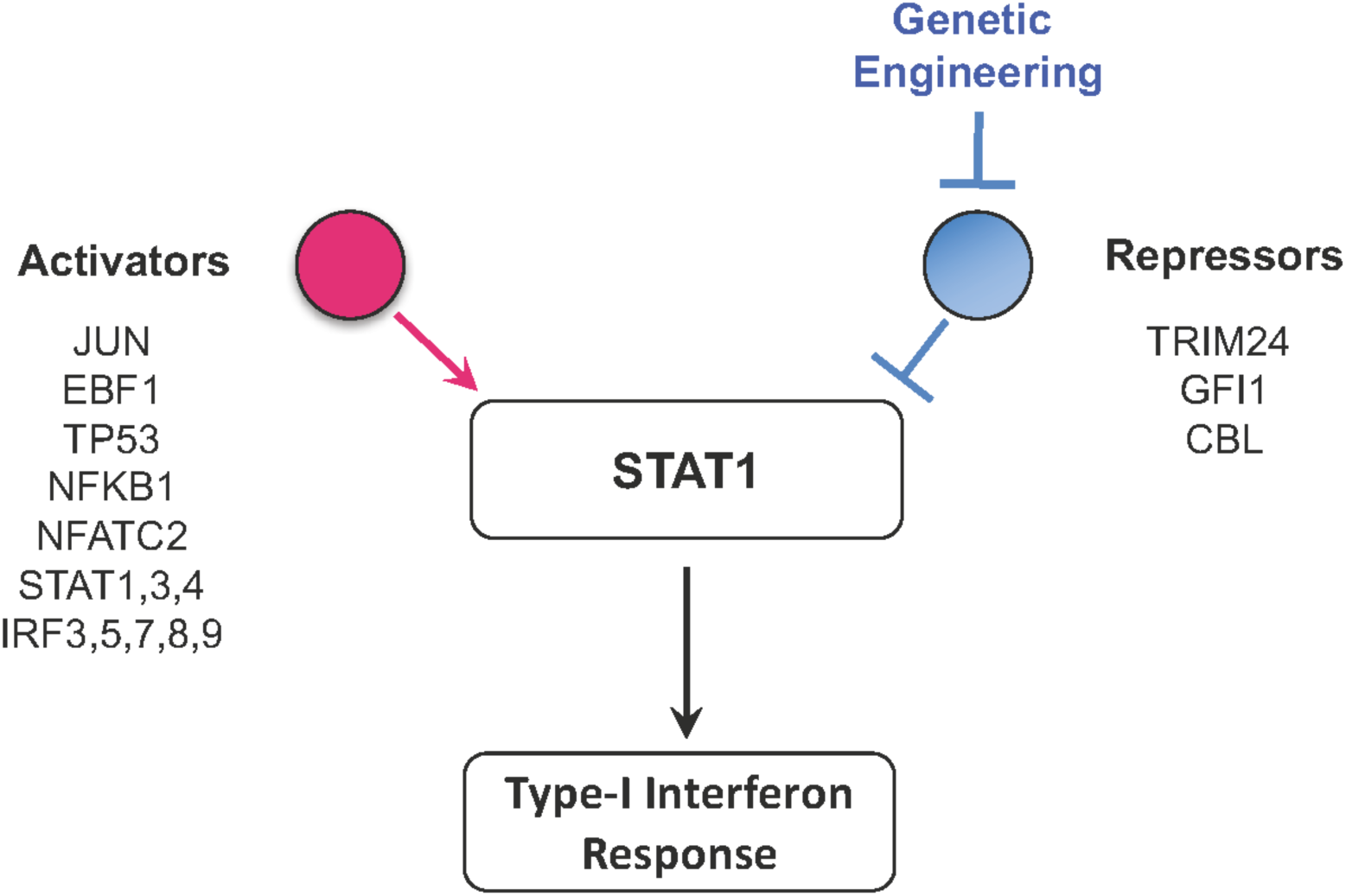
Identification of regulators of STAT1 as candidates for engineering the antiviral response. Schematic of the regulators of STAT1, which may be candidates for engineering and improving virus resistance in CHO cells.

**Figure 6.**
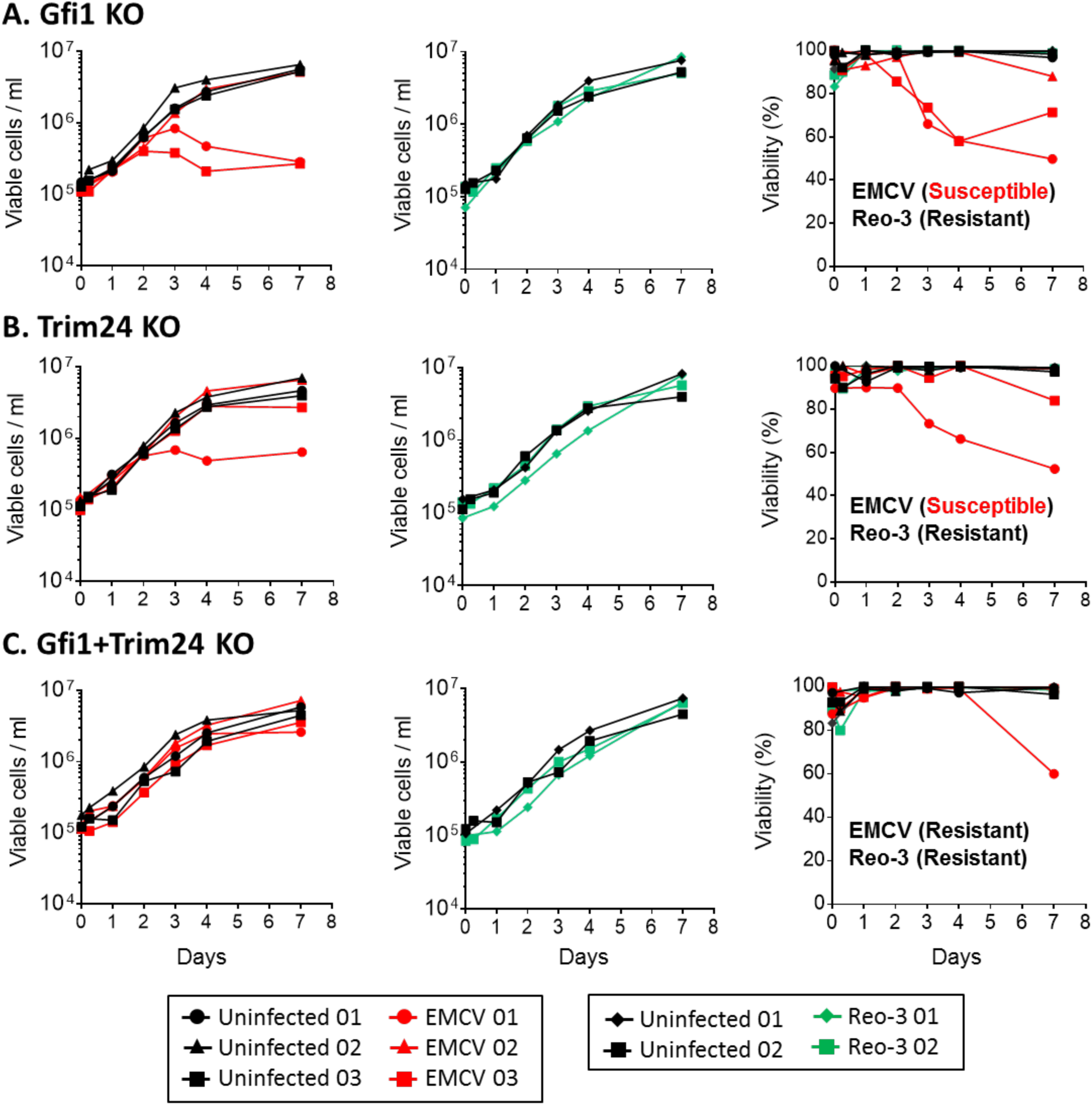
Viral resistance of the Gfi1 and/or Trim24 KO engineered CHO cells. Gfi1 and Trim24 were knocked out and tested for resistance to EMCV and Reo-3 virus infection compared to the control (susceptible) cells. Cell density and viability was followed up for one week post infection (p.i.) for Gfi1 single knockout cells (A), Trim24 single knockout cells (B) and Gfi1 and Trim24 double knockout cells (C). Data shown is from three (EMCV) and two (Reo-3) independent virus infection experiments. Susceptible CHO cell lines were used as positive controls for EMCV and Reo-3 virus infections during the first seven days (Figure S10). In some experiments, resistant cultures were passaged and followed up for an additional week (Figure S11).

## Conclusions

Here we perform a genome-wide study of viral resistance in CHO, thereby demonstrating the utility of systems biology approaches to not only improve host cell productivity and metabolism^70-72^, but also to improve product safety. Specifically, we demonstrated that STAT1 and other key regulators contribute to the inhibition of RNA virus replication in CHO cell lines. Furthermore, an analysis of poly I:C treatment exposed these molecular mechanisms underlie the protection against RNA virus infection. Studies have shown that modulating genetic factors can promote viral resistance in CHO cells^11, 73, 74^. However, our findings suggest novel cell engineering targets beyond those coding for cell receptors. Thus, these insights provide further tools to enable the development of virus-resistant hosts to improve safety and secure the availability of biotherapeutic products^3, 75, 76^.

## Acknowledgments

This work was supported by generous funding from the Novo Nordisk Foundation provided to the Center for Biosustainability at the Technical University of Denmark (grant no. NNF10CC1016517). Funding was also received from the Regulatory Science and Review Enhancement program at the FDA (RSR #14-07).

## Author contributions

ASR conceived of the project idea; MP, ASR, NEL directed the research; AWTC, MP, and NEL wrote the manuscript; AWTC, SL, BPK, CCK, JMG, and FG analyzed the RNA-Seq data; PM, SPB, BGV conducted the RNA-Seq; MP, GC, YZ and HS conducted CHO cell and virus experiments. All authors read and approved of this work.

## Reference

1. Walsh, G. Biopharmaceutical benchmarks 2014. Nat Biotechnol 32, 992-1000 (2014).

2. Weiebe, M.E. et al. A multifaceted approach to assure that recombinant tPA is free of adventitious virus. In: Advances in animal cell biology and technology for bioprocesses. (Spier, Griffitlis, Stephenne, Crooy, eds.). 68-71 (1989).

3. Berting, A., Farcet, M.R. & Kreil, T.R. Virus susceptibility of Chinese hamster ovary (CHO) cells and detection of viral contaminations by adventitious agent testing. Biotechnol Bioeng 106, 598-607 (2010).

4. Poiley, J.A., Nelson, R.E., Hillesund, T. & Rainer, i.R. Susceptibility of cho k1 cells to infection by eight adventitious viruses and four retroviruses. In Vitro Toxicology 4, 1-12 (1991).

5. Garnick, R.L. Raw materials as a source of contamination in large-scale cell culture. Dev Biol Stand 93, 21-29 (1998).

6. Nims, R.W. Detection of adventitious viruses in biologicals‐‐a rare occurrence. Dev Biol (Basel) 123, 153-164; discussion 183-197 (2006).

7. Dinowitz, M. et al. Recent studies on retrovirus-like particles in Chinese hamster ovary cells. Dev Biol Stand 76, 201-207 (1992).

8. Rabenau, H. et al. Contamination of genetically engineered CHO-cells by epizootic haemorrhagic disease virus (EHDV). Biologicals 21, 207-214 (1993).

9. Bethencourt, V. Virus stalls Genzyme plant. Nature Biotechnology 27, 681 (2009).

10. Merten, O.W. Virus contaminations of cell cultures - A biotechnological view. Cytotechnology 39, 91-116 (2002).

11. Mascarenhas, J.X. et al. Genetic engineering of CHO cells for viral resistance to minute virus of mice. Biotechnol Bioeng 114, 576-588 (2017).

12. Schoggins, J.W. & Rice, C.M. Interferon-stimulated genes and their antiviral effector functions. Curr Opin Virol 1, 519-525 (2011).

13. Sadler, A.J. & Williams, B.R. Interferon-inducible antiviral effectors. Nat Rev Immunol 8, 559-568 (2008).

14. Perry, A.K., Chen, G., Zheng, D., Tang, H. & Cheng, G. The host type I interferon response to viral and bacterial infections. Cell Res 15, 407-422 (2005).

15. Taniguchi, T. & Takaoka, A. The interferon-alpha/beta system in antiviral responses: a multimodal machinery of gene regulation by the IRF family of transcription factors. Curr Opin Immunol 14, 111-116 (2002).

16. Pantelic, L., Sivakumaran, H. & Urosevic, N. Differential induction of antiviral effects against West Nile virus in primary mouse macrophages derived from flavivirus-susceptible and congenic resistant mice by alpha/beta interferon and poly(I-C). J Virol 79, 1753-1764 (2005).

17. Green, T.J. & Montagnani, C. Poly I:C induces a protective antiviral immune response in the Pacific oyster (Crassostrea gigas) against subsequent challenge with Ostreid herpesvirus (OsHV-1 muvar). Fish Shellfish Immunol 35, 382-388 (2013).

18. Plant, K.P., Harbottle, H. & Thune, R.L. Poly I:C induces an antiviral state against Ictalurid Herpesvirus 1 and Mx1 transcription in the channel catfish (Ictalurus punctatus). Dev Comp Immunol 29, 627-635 (2005).

19. Lewis, N.E. et al. Genomic landscapes of Chinese hamster ovary cell lines as revealed by the Cricetulus griseus draft genome. Nature Biotechnology 31, 759-+ (2013).

20. Xu, X. et al. The genomic sequence of the Chinese hamster ovary (CHO)-K1 cell line. Nat Biotechnol 29, 735-741 (2011).

21. Vishwanathan, N. et al. Augmenting Chinese hamster genome assembly by identifying regions of high confidence. Biotechnol J 11, 1151-1157 (2016).

22. Chen, C., Le, H. & Goudar, C.T. Evaluation of two public genome references for Chinese hamster ovary cells in the context of RNA-seq based gene expression analysis. Biotechnol Bioeng (2017).

23. van Wijk, X.M. et al. Whole-Genome Sequencing of Invasion-Resistant Cells Identifies Laminin alpha2 as a Host Factor for Bacterial Invasion. MBio 8 (2017).

24. Wang, Z., Gerstein, M. & Snyder, M. RNA-Seq: a revolutionary tool for transcriptomics. Nat Rev Genet 10, 57-63 (2009).

25. Vishwanathan, N. et al. Global Insights Into the Chinese Hamster and CHO Cell Transcriptomes. Biotechnology and Bioengineering 112, 965-976 (2015).

26. Yuk, I.H. et al. Effects of copper on CHO cells: insights from gene expression analyses. Biotechnol Prog 30, 429-442 (2014).

27. Fomina-Yadlin, D. et al. Transcriptome analysis of a CHO cell line expressing a recombinant therapeutic protein treated with inducers of protein expression. J Biotechnol 212, 106-115 (2015).

28. Hsu, H.H. et al. A Systematic Approach to Time-series Metabolite Profiling and RNA-seq Analysis of Chinese Hamster Ovary Cell Culture. Sci Rep 7, 43518 (2017).

29. Puig, M. et al. TLR9 and TLR7 agonists mediate distinct type I IFN responses in humans and nonhuman primates in vitro and in vivo. J Leukocyte Biol 91, 147-158 (2012).

30. Bolger, A.M., Lohse, M. & Usadel, B. Trimmomatic: a flexible trimmer for Illumina sequence data. Bioinformatics 30, 2114-2120 (2014).

31. Dobin, A. et al. STAR: ultrafast universal RNA-seq aligner. Bioinformatics 29, 15-21 (2013).

32. Anders, S., Pyl, P.T. & Huber, W. HTSeq‐‐a Python framework to work with high-throughput sequencing data. Bioinformatics 31, 166-169 (2015).

33. Anders, S. & Huber, W. Differential expression analysis for sequence count data. Genome Biol 11, R106 (2010).

34. Katze, M.G., He, Y. & Gale, M., Jr. Viruses and interferon: a fight for supremacy. Nat Rev Immunol 2, 675-687 (2002).

35. Seo, Y.J. & Hahm, B. Type I interferon modulates the battle of host immune system against viruses. Adv Appl Microbiol 73, 83-101 (2010).

36. McNab, F., Mayer-Barber, K., Sher, A., Wack, A. & O’Garra, A. Type I interferons in infectious disease. Nat Rev Immunol 15, 87-103 (2015).

37. Schneider, W.M., Chevillotte, M.D. & Rice, C.M. Interferon-Stimulated Genes: A Complex Web of Host Defenses. Annu Rev Immunol 32, 513-545 (2014).

38. Ahmed, M. et al. Ability of the matrix protein of vesicular stomatitis virus to suppress beta interferon gene expression is genetically correlated with the inhibition of host RNA and protein synthesis. J Virol 77, 4646-4657 (2003).

39. Rieder, M. & Conzelmann, K.K. Rhabdovirus Evasion of the Interferon System. J Interf Cytok Res 29, 499-509 (2009).

40. Sherry, B. Rotavirus and Reovirus Modulation of the Interferon Response. J Interf Cytok Res 29, 559-567 (2009).

41. Ng, C.S. et al. Encephalomyocarditis Virus Disrupts Stress Granules, the Critical Platform for Triggering Antiviral Innate Immune Responses. J Virol 87, 9511-9522 (2013).

42. Jensen, S. & Thomsen, A.R. Sensing of RNA viruses: a review of innate immune receptors involved in recognizing RNA virus invasion. J Virol 86, 2900-2910 (2012).

43. Goubau, D. et al. Antiviral immunity via RIG-I-mediated recognition of RNA bearing 5’ diphosphates. Nature 514, 372-+ (2014).

44. Loo, Y.M. et al. Distinct RIG-I and MDA5 signaling by RNA viruses in innate immunity. J Virol 82, 335-345 (2008).

45. Honda, K. & Taniguchi, T. IRFs: master regulators of signalling by Toll-like receptors and cytosolic pattern-recognition receptors. Nat Rev Immunol 6, 644-658 (2006).

46. Ivashkiv, L.B. & Donlin, L.T. Regulation of type I interferon responses. Nat Rev Immunol 14, 36-49 (2014).

47. Jowett, J.B. et al. The human immunodeficiency virus type 1 vpr gene arrests infected T cells in the G2 + M phase of the cell cycle. J Virol 69, 6304-6313 (1995).

48. Krauer, K.G. et al. The Epstein-Barr virus nuclear antigen-6 protein co-localizes with EBNA-3 and survival of motor neurons protein. Virology 318, 280-294 (2004).

49. Darbinyan, A. et al. Evidence for dysregulation of cell cycle by human polyomavirus, JCV, late auxiliary protein. Oncogene 21, 5574-5581 (2002).

50. Everett, R.D., Earnshaw, W.C., Findlay, J. & Lomonte, P. Specific destruction of kinetochore protein CENP-C and disruption of cell division by herpes simplex virus immediate-early protein Vmw110. EMBO J 18, 1526-1538 (1999).

51. Kramer, A., Green, J., Pollard, J., Jr. & Tugendreich, S. Causal analysis approaches in Ingenuity Pathway Analysis. Bioinformatics 30, 523-530 (2014).

52. Olive, C. Pattern recognition receptors: sentinels in innate immunity and targets of new vaccine adjuvants. Expert Rev Vaccines 11, 237-256 (2012).

53. Bohlson, S.S. Modulators of the innate immune response. Curr Drug Targets 9, 101 (2008).

54. Mutwiri, G., Gerdts, V., Lopez, M. & Babiuk, L.A. Innate immunity and new adjuvants. Rev Sci Tech 26, 147-156 (2007).

55. Diamond, M.S. & Farzan, M. The broad-spectrum antiviral functions of IFIT and IFITM proteins. Nat Rev Immunol 13, 46-57 (2013).

56. Li, K. et al. IFITM proteins restrict viral membrane hemifusion. PLoS Pathog 9, e1003124 (2013).

57. Pillai, P.S. et al. Mx1 reveals innate pathways to antiviral resistance and lethal influenza disease. Science 352, 463-466 (2016).

58. Verhelst, J., Hulpiau, P. & Saelens, X. Mx proteins: antiviral gatekeepers that restrain the uninvited. Microbiol Mol Biol Rev 77, 551-566 (2013).

59. Walsh, D., Mathews, M.B. & Mohr, I. Tinkering with translation: protein synthesis in virus-infected cells. Cold Spring Harb Perspect Biol 5, a012351 (2013).

60. Arpaia, N. & Barton, G.M. Toll-like receptors: key players in antiviral immunity. Curr Opin Virol 1, 447-454 (2011).

61. Kawai, T. & Akira, S. The roles of TLRs, RLRs and NLRs in pathogen recognition. Int Immunol 21, 317-337 (2009).

62. Thompson, A.J. & Locarnini, S.A. Toll-like receptors, RIG-I-like RNA helicases and the antiviral innate immune response. Immunol Cell Biol 85, 435-445 (2007).

63. Aaronson, D.S. & Horvath, C.M. A road map for those who don’t know JAK-STAT. Science 296, 1653-1655 (2002).

64. Au-Yeung, N., Mandhana, R. & Horvath, C.M. Transcriptional regulation by STAT1 and STAT2 in the interferon JAK-STAT pathway. JAKSTAT 2, e23931 (2013).

65. Li, H.S. & Watowich, S.S. Innate immune regulation by STAT-mediated transcriptional mechanisms. Immunol Rev 261, 84-101 (2014).

66. Hu, X. & Ivashkiv, L.B. Cross-regulation of signaling pathways by interferon-gamma: implications for immune responses and autoimmune diseases. Immunity 31, 539-550 (2009).

67. Fortier, M.E. et al. The viral mimic, polyinosinic:polycytidylic acid, induces fever in rats via an interleukin-1-dependent mechanism. Am J Physiol Regul Integr Comp Physiol 287, R759-766 (2004).

68. Sharif-Askari, E. et al. Zinc finger protein Gfi1 controls the endotoxin-mediated Toll-like receptor inflammatory response by antagonizing NF-kappaB p65. Mol Cell Biol 30, 3929-3942 (2010).

69. Tisserand, J. et al. Tripartite motif 24 (Trim24/Tif1alpha) tumor suppressor protein is a novel negative regulator of interferon (IFN)/signal transducers and activators of transcription (STAT) signaling pathway acting through retinoic acid receptor alpha (Raralpha) inhibition. J Biol Chem 286, 33369-33379 (2011).

70. Gutierrez, J.M. & Lewis, N.E. Optimizing eukaryotic cell hosts for protein production through systems biotechnology and genome-scale modeling. Biotechnol J 10, 939-949 (2015).

71. Kuo, C.C. et al. The emerging role of systems biology for engineering protein production in CHO cells. Curr Opin Biotechnol 51, 64-69 (2018).

72. Richelle, A. & Lewis, N.E. Improvements in protein production in mammalian cells from targeted metabolic engineering. Curr Opin Syst Biol 6, 1-6 (2017).

73. Taber, R., Alexander, V. & Wald, N., Jr. The selection of virus-resistant Chinese hamster ovary cells. Cell 8, 529-533 (1976).

74. Haines, K.M., Vande Burgt, N.H., Francica, J.R., Kaletsky, R.L. & Bates, P. Chinese hamster ovary cell lines selected for resistance to ebolavirus glycoprotein mediated infection are defective for NPC1 expression. Virology 432, 20-28 (2012).

75. FDA Guidance for industry: Q5A viral safety evaluation of biotechnology products derived from cell lines of human or animal origin. (1998).

76. FDA Guidance for industry: Characterization and qualification of cell substrates and other biological starting materials used in the production of viral vaccines for the prevention and treatment of infectious diseases. (2006).

